# Integrative Network Analysis of Differentially Methylated and Expressed Genes for Biomarker Identification in Leukemia

**DOI:** 10.1101/658948

**Authors:** Robersy Sanchez, Sally A. Mackenzie

## Abstract

Genome-wide DNA methylation and gene expression are commonly altered in pediatric acute lymphoblastic leukemia (PALL). Integrated analysis of cytosine methylation and expression datasets has the potential to provide deeper insights into the complex disease states and their causes than individual disconnected analyses. Studies of protein-protein interaction (PPI) networks of differentially methylated (DMGs) and expressed genes (DEGs) showed that gene expression and methylation consistently targeted the same gene pathways associated with cancer: *Pathways in cancer, Ras signaling pathway, PI3K-Akt signaling pathway*, and *Rap1 signaling pathway*, among others. Detected gene hubs and hub sub-networks are integrated by signature loci associated with cancer that include, for example, *NOTCH1, RAC1, PIK3CD, BCL2*, and *EGFR*. Statistical analysis disclosed a stochastic deterministic dependence between methylation and gene expression within the set of genes simultaneously identified as DEGs and DMGs, where larger values of gene expression changes are probabilistically associated with larger values of methylation changes. Concordance analysis of the overlap between enriched pathways in DEG and DMG datasets revealed statistically significant agreement between gene expression and methylation changes, reflecting a coordinated response of methylation and gene-expression regulatory systems. These results support the identification of reliable and stable biomarkers for cancer diagnosis and prognosis.

## Introduction

Network-based modeling approaches have the potential to integrate and improve the perception of complex disease states and their root causes. To date, network analysis provides reliable and cost effective approaches for early disease detection, prediction of co-occurring diseases and interactions, and drug design ^1^. Although integrated genomic analysis of methylation and gene expression in leukemia have been performed ^2–5^, an integration including network analysis of methylation and gene expression is still missing.

Our study investigates protein-protein interaction networks (PPI), which are exclusively focused on protein-protein associations and resulting cell activities. A PPI network can be defined as a (un)directed graph/network holding vertices as proteins (or protein-encoding genes) and edges as the interactions/association between them. Associations are meant to be specific and biologically meaningful, i.e., two proteins (genes) are connected by an edge if jointly contributing to a shared function, which does not necessarily reflect a physical binding interaction.

Within the network, some proteins denote hubs interacting with numerous partners. Biologically, hubs are key elements on which functionality of the cellular process modeled by the network depends. Consequently, it is reasonable to assume that a biomarker suitable to define specific disease states would likely be a hub or a hub regulator within a relevant network. Frequently, more than one interacting network model is possible, with each model carrying a different uncertainty level for the biological process under study. Integration of more than one network model can help to reduce the implicit uncertainty associated to each model prediction^6^.

Here, we address the hypothesis that disease-induced DNA methylation changes can serve as a source of reliable and stable biomarkers for cancer diagnosis and prognosis. Toward that aim, aberrant DNA methylation of key genes was reported in Acute Lymphoblastic Leukemia (ALL) ^6^. We report on a reproducible approach integrating network analysis of DMGs, DEGs and DEGs-DMGs estimated within datasets from patients with pediatric ALL (PALL). Such an integration may provide the basis for robust identification of reliable and stable biomarkers for cancer diagnosis and prognosis.

## Results

### General methylation features of the study

Differentially methylated positions (DMPs) were estimated for control (four normal CD19+ blood cell donors) and patient (ALL cells from three patients) groups relative to a reference group of four independent normal CD19+ blood cell donors. Inclusion of a reference group permitted the evaluation of natural variation in healthy individuals and reduction of noise in a signal detection step of our methylation analysis pipeline. The distribution of methylation changes at DMPs along the chromosome revealed a genome-wide methylation re-patterning dominated by hypermethylation in PALL patients (Supplementary Fig. S1). Hypomethylated sites are visible in the genome browser after zooming (tracks available in the Supplementary File S1). Consistent with natural methylation variability in the population of healthy individuals, DMPs were observed in the control group as well.

DMGs were estimated from group comparisons for number of DMPs within gene-body regions between control (CD19+ blood cell donors) and ALL cells based on generalized linear regression. This analysis yielded a total of 4795 DMGs, including protein-coding regions (3338) and non-coding RNA genes (Supplementary Table S1). 1774 genes from the set of 2360 reported (B-Cells) DEGs in the original study ^7^ were DMGs as well (75.2%, Supplementary Table S2).

The gene-body methylation signal detected in PALL patients coincided with a significant number of genes from the list of all cancer consensus genes (723) from the COSMIC database^8^: 254 DMGs, and 126 DEGs, and from them 112 DEGs-DMGs.

### Network analysis on a set of differentially methylated genes (DMGs)

When applying network analysis, not all DMGs and DEGs estimated from the experimental datasets integrate networks. A subset of the most relevant genes from the experimental dataset able to integrate networks is helpful to facilitate further network analysis. The preliminary application of network-based enrichment analysis (NBEA^9^) and network enrichment analysis test (NEAT ^10^) on the set of DMGs permitted the identification of 285 network-related DMGs (Supplementary Tables S1 and S1). Similar analysis permitted the identification of 326 network-related DEGs (Supplementary Table S2-B, from B-Cells 2360 DEGs reported in Supplementary Table 3 from original study ^7^). These subsets were used to build the corresponding protein-protein interaction (PPI) networks with the STRING app of Cytoscape ^11,12^. Alternatively, to bypass possible bias introduced by the heuristic used to subset the whole set of genes (NBEA^9^ and NEAT^10^), sub-clusters of hubs where retrieved applying the MCODE Cytoscape app on the whole set of DMGs.

The PPI network built on the set of 285 DMGs is presented in Supplementary Fig. S2. The analysis with available tools in Cytoscape ^11^ led to the identification of the main hubs from the PPI network (Fig. 1A and C). Sizes of nodes and labels, as well as their colors, are used for rapid identification of network hubs. Network hubs were confirmed based on *betweenness-centrality* and *node degree*^13^, such that the size of each node is proportional to its value of *betweenness-centrality* and the label font size is proportional to its *node degree*.

**Figure 1.**
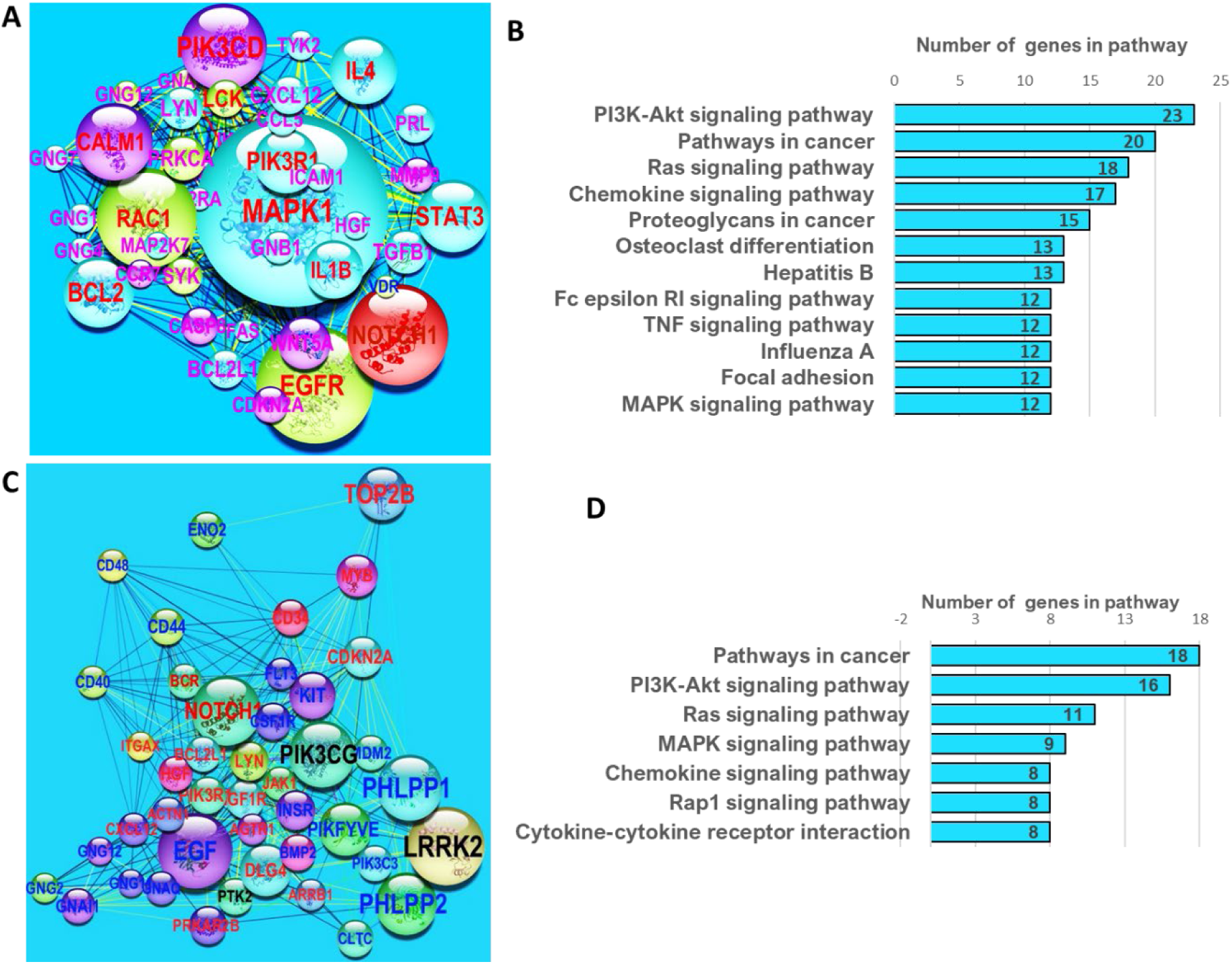
PPI subnetworks of hubs derived from subsets of network-related DMGs. **A**, main subnetwork of hubs obtained with the application of K-means clustering on the set of 285 network-related DMGs identified with NBEA^9^ and NEAT^10^ tests. The size of each node is proportional to its value of betweeness centrality and the label font size is proportional to its node degree. Node colors from light-green to red maps the discrete scale of logarithm base 2 of fold changes in DMP numbers for the corresponding gene: light-green: [1, 2), cyan: [2, 3), blue: [3, 4), and red: 5 or more. Edge color is based on co-expression index from white (0.042) to red (0.842). **B**, enrichment analysis with Cytoscape11 on KEGG pathway sets on network in **A. C**, main subnetwork of hubs obtained with the application of MCODE Cytoscape app and K-means clustering. **D**, enrichment analysis with Cytoscape11 on KEGG pathway sets on the network in **C**.

The main hub subnetworks in Fig. 1A and 1C were identified with the application of K-means clustering on the main networks shown in Supplementary Fig. S2 and S3, respectively, with network centralities measuring *Degree, Betweeness-Centrality, Closeness-Centrality, Clustering-Coefficient*, and *Average-Shortest-Path*. Network enrichment analysis of the subnetwork of hubs identified KEGG pathways involved in cancer development (Fig. 1B and D), supporting our findings with analysis of network centralities.

K-means clustering split the network of 285 DMGs (Supplementary Fig. S2) into three clusters: *i*) the main subnetwork of hubs (46 DMGs, shown in Fig.1A, Supplementary Table S1), *ii*) a subnetwork with minor hubs (101 DMGs, Supplementary Fig. S4, Supplementary Table S1), and *iii*) a cluster integrated by two subnetworks (139 DMGs, Supplementary Fig. S4, Supplementary Table S1). Results with MCODE Cytoscape app and K-means were consistent with those obtained by subsetting the whole set of DMGs via NBEA and NEAT ^9,10^ (Supplementary Fig. S3 and Supplementary Tables S2), with a notable enrichment of KEGG pathways associated with cancer development (Supplementary Fig. S5).

The scatter plots of network centrality measures (Fig. 2) suggest that the main subnetwork of hubs includes the most relevant network nodes/genes (in red) carrying the highest network centrality measurements. We noted a transition from a non-linear behavior, in clusters *iii* (nodes in blue) and *ii* (node in green), to a linear trend observed in cluster *i* (red points, Fig. 2). These analyses suggest that the subnetwork of hubs shown in Fig. 1C also involves genes with methylation signals that have a role in PALL development ^14^.

**Figure 2.**
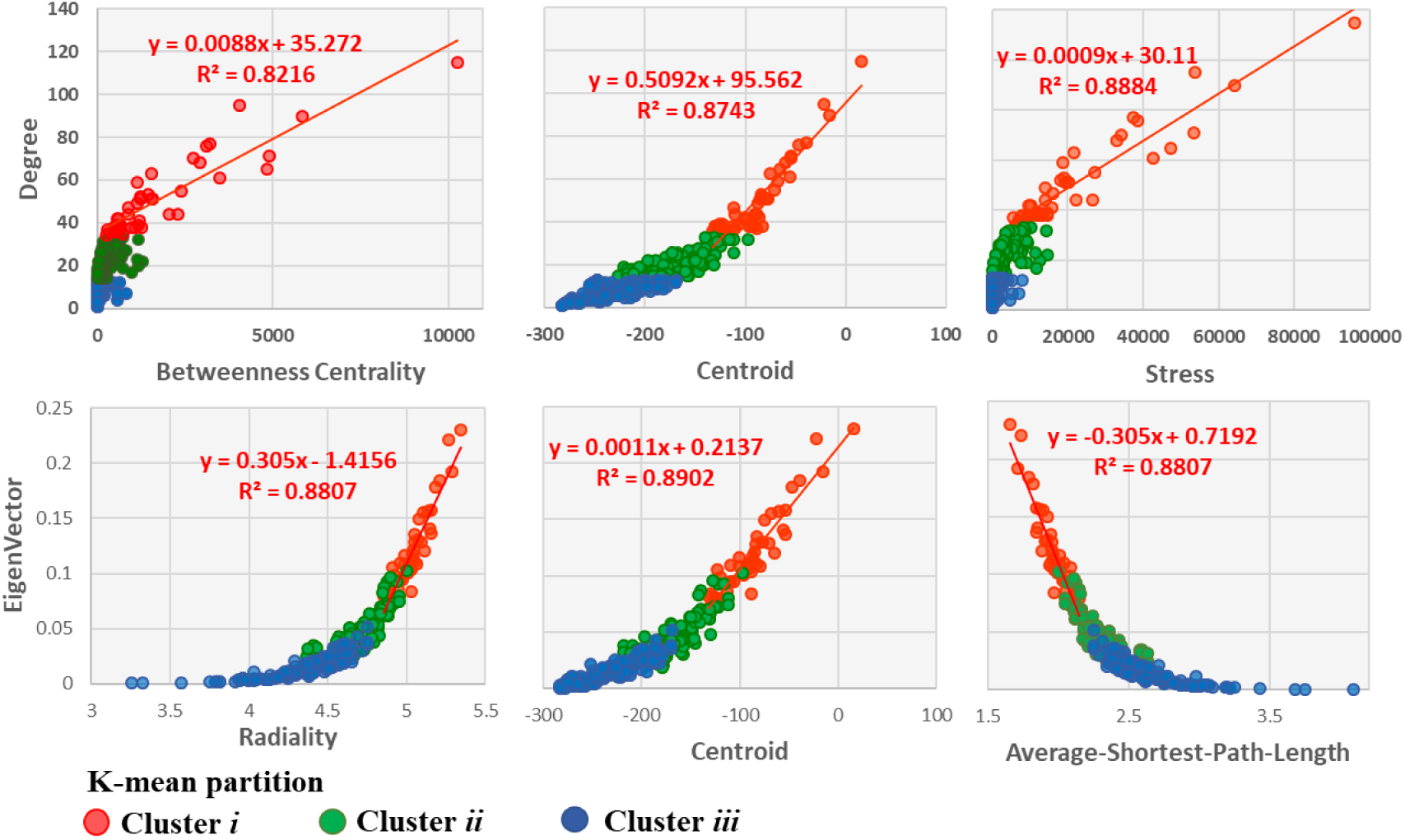
Scatter plots of network centralities measures. A general non-linear trend is notable for genes/nodes from clusters *iii* to *ii*, while the linear trend in cluster *i* can be visualized. The highest values of network centralities: *degree, betweenness, centroid, stress*, and *radiality*, are found in cluster *i*, which correspond to the main subnetwork of hubs presented in Fig. 1B (consistent with the lowest values of *average-shortest-path-length*). Networks from clusters *i, ii*, and *iii* are shown in Supplementary Fig. S4.

Results of network enrichment analysis of DMG and DEG PPI networks built with STRING (Cytoscape) are shown in Fig. 3 (Supplementary Tables S1 an S2). The analyses indicate that DMG and DEG datasets targeted many of the same pathways with overlap of 80% (Fig. 3C). Pathways linked to cancer development and apoptosis are notable, and KEGG *pathways in cancer* (hsa05200) showed pronounced enrichment, with more than 50 and 40 genes from the DMG and DEG datasets, respectively.

**Figure 3.**
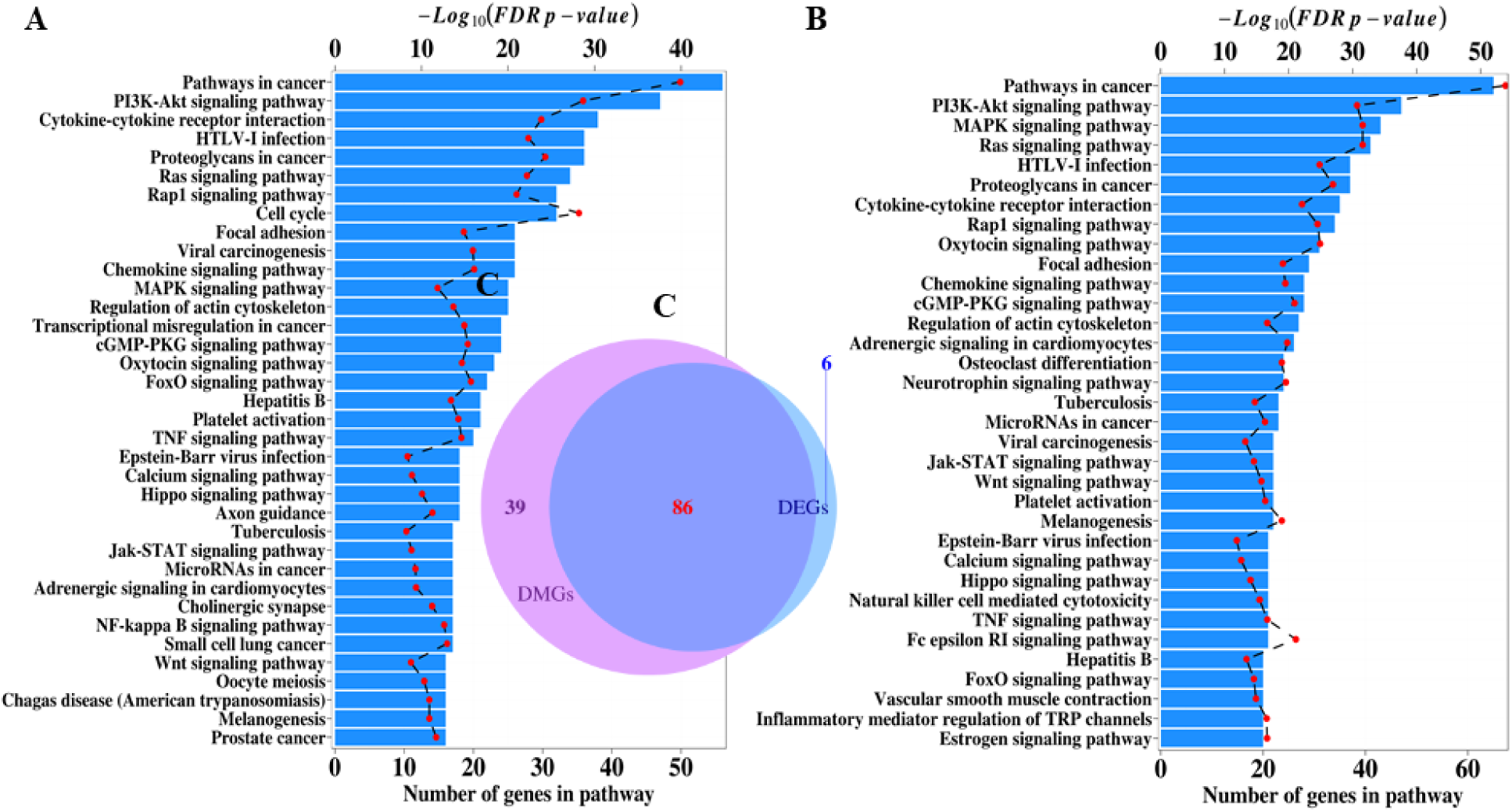
Network based enrichment analysis of protein-protein interaction (PPI) networks independently derived from DMGs and DEGs estimated in patients with PALL. **A**, PPI enriched network of DEGs with 15 or more genes. **B**, PPI enriched network of DMGs with 20 or more genes. **C**, Venn diagram with the overlapping of all PPI enriched networks of DMGs and DEGs with 7 or more genes. The PPI enriched network analysis was performed in STRING app on Cytoscape, 11,12 and the analysis is limited to KEGG human pathways.

In the case of patients with PALL, enrichment for *PI3K-Akt signaling pathway, MAPK signaling pathway, JAK-STAT signaling pathway, Wnt signaling pathway*, and *Focal adhesion* (all included in KEGG *pathway in cancer*) was statistically significant for both DMG and DEG subsets. The Venn diagram shown in Fig. 3C implies a high level of concordance between the enriched KEGG pathways identified in PPI networks from DEGs and from DMGs.

Figure 4 supports a strong concordance between the enriched KEGG pathways identified in PPI networks from DEGs and from DMGs. Bootstrap Bayesian estimation of the Lin’s concordance correlation coefficient (*ρ*_*CC*_) yielded a value of *ρ*_*CC*_= 0.71 with a confidence interval (C.I.) 0.52 ≤ *ρ*_*CC*_≤ 0.84, and a Kendall coefficient of concordance *ρ*_*KC*_= 081 (permutation *p*-value < 0.001). The linear regression analysis presented in Fig. 4A indicates a statistically significant linear relationship between the *pathway score* (*P*_*DMG*_) of enriched KEGG pathways in DMG PPI network (see definition at equation (1)) and *pathway score* (*P*_*DMG*_) of enriched KEGG pathways in DMG PPI network. The proximity of most of the regression points (pairs of pathways scores) around the identity line (dashed line in blue) suggests significant agreement between methylation and gene expression regulatory systems, also indicated by aregression slope of 0.9. This concordance between gene expression and methylation is graphically corroborated by Bland-Altman plot ^15^, where almost all the points are located in between the *mean* − 2σ and *mean* + 2σ horizontal lines (Fig. 4B).

**Figure 4.**
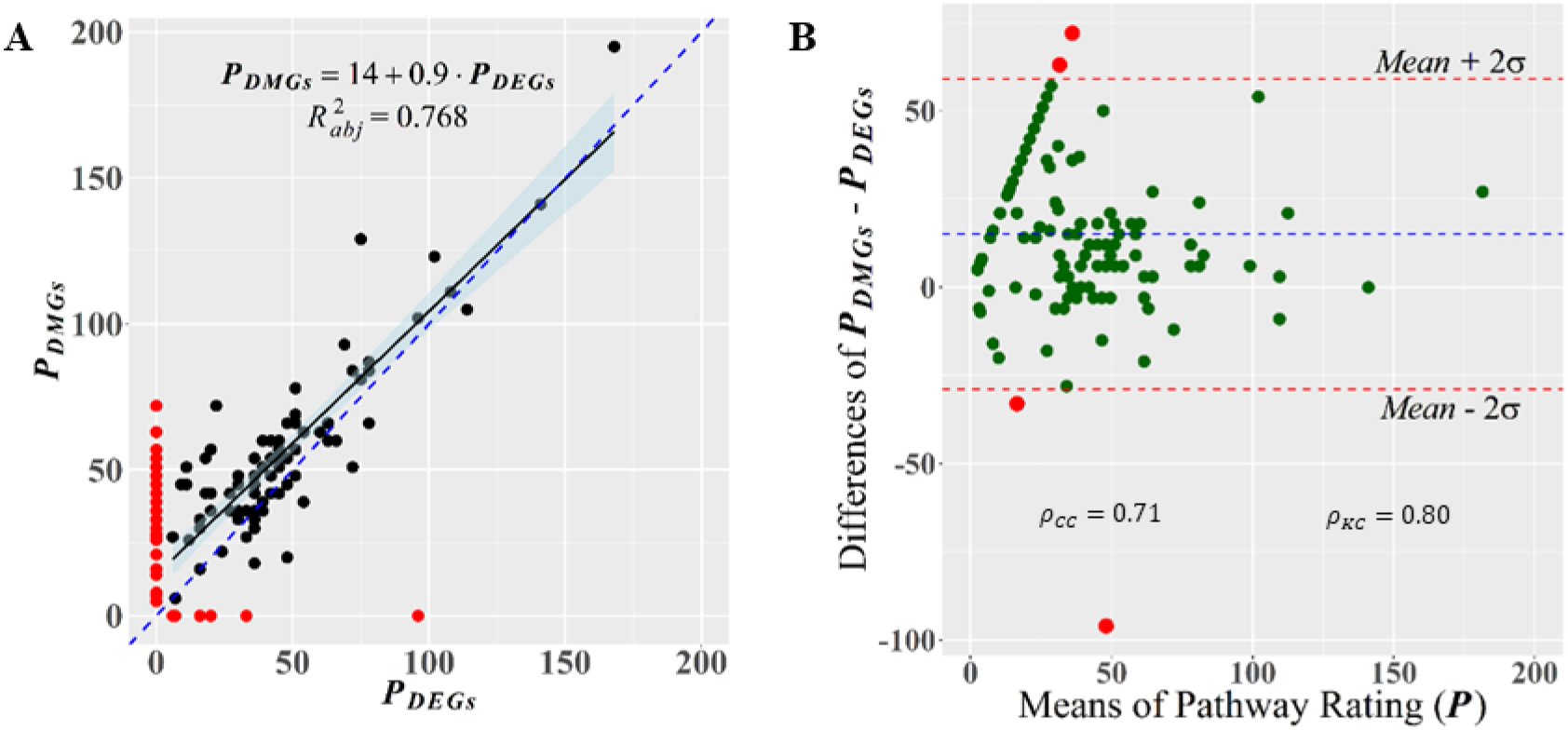
Graphical evaluation of the concordance between DEG and DMG enrichments on KEGG pathways. **A**, scatterplot of pathway ratings (see Eq. 1) from enriched pathways on the set of DMGs (*P*_*DMGS*_) and DEGs (*P*_*DMGS*_), respectively. The regression analysis shows the linear trend of the relationship *P*_*DMGS*_ > 0 *versus P_DMGS_* > 0 (black dots). The identity dashed line (in blue) helps in gauging the degree of agreement between measurements ^15^. Dots in red highlight pathways for which *P*_*DMGS*_ = 0 or *P*_*DMGS*_ = 0. **B**, Bland-Altman plot of the agreement, on targeting gene pathways, between responses from gene expression and methylation regulatory systems. The agreement between measurements can also be tested by values of the Lin’s concordance correlation coefficient (*ρ*_*CC*_) and Kendall coefficient of concordance (*ρ* _*KC*_).

### DEG-DMG network analysis

NBEA and NEAT ^9,10^ were applied to identify network-related genes from the set of DEGs-DMGs (191, 1774 genes). The PPI network of 191 DEGs-DMGs is shown in Supplementary Fig. S6 (Supplementary Table S2). Three clusters were detected by applying K-means clustering on the main PPI-network of DEGs-DMGs and two of them integrated the subnetworks of hubs shown in Fig. 5B and D, while the third cluster gave rise to several subsets of subnetworks.

**Figure 5.**
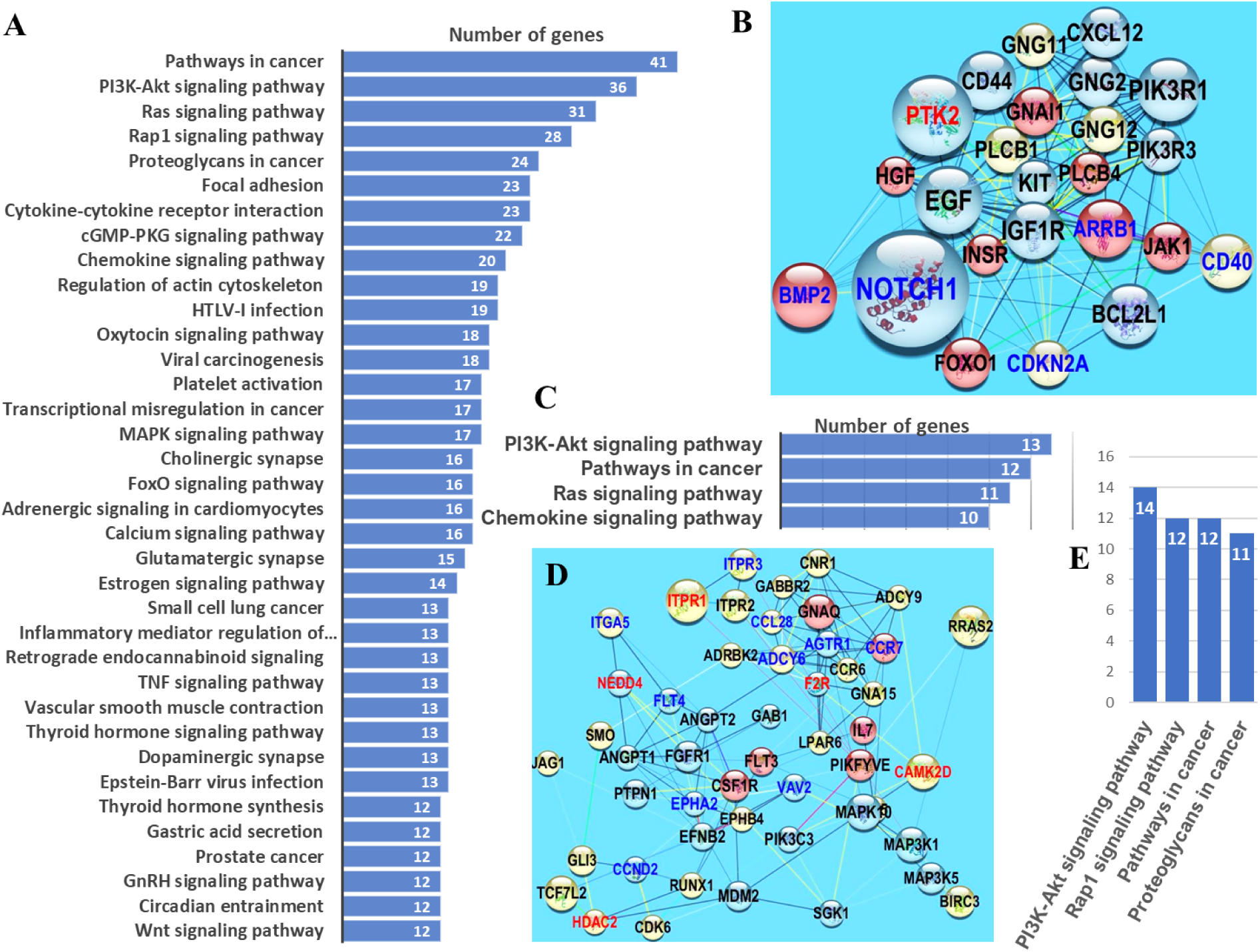
Network enrichment for the network-related DEG-DMGs. **A**, bar-plots of the enriched KEGG pathways in the PPI-network of 191 DEG-DMGs (Supplementary Fig. S6). **B** and **D**, subnetworks integrated by gene-hubs identified with K-means clustering of the network from panel. **C** and **E**, bar-plots of the enriched KEGG pathways on the networks from panels B and D, respectively. In the networks, nodes with the same color belong to the same cluster obtained with K-Medoid clustering. Gene hubs were identified based on *betweeness centrality* and *node degree*, such that the size of each node is proportional to its value of *betweeness centrality* and the label font size is proportional to its *node degree*. Edge color is based on *coexpression* index from white (0.042) to red (0.938). The PPI network and the enrichment analyses were performed in STRING app on Cytoscape ^11,12^.

Enrichments detected in the main PPI network of 191 DEGs-DMGs network (Fig. 5A) and subnetworks (Fig. 5C and 5E. Supplementary Table S2)) are consistent with previous results (Fig. 3): *i*) only focused on the set of DMGs (not all of them DEGs, Fig. 3A) and *ii*) only focused on the set of DEGs (not all of them DMGs, Fig. 3B).

Group means of methylation level differences at each gene-body DMP for genes NOTCH1, CD44, and BCL2L1 (hubs from the DMGs-DEGs sub-network from Fig. 5B) are shown in Fig. 6A. NOTCH1 and CD44 have been reported to be epigenetically regulated ^16–19^ and, in particular, NOTCH1 has been proposed as a drug target for the treatment of T-cell acute lymphoblastic leukemia ^17^. BCL2L1 is known to have roles in apoptosis and has been proposed as a drug target for cancer treatment ^20^. Genes from activation of the mitogen-activated protein kinase (MAPK) pathway are frequently altered in cancer and have been proposed as drug targets as well ^21^.

**Figure 6.**
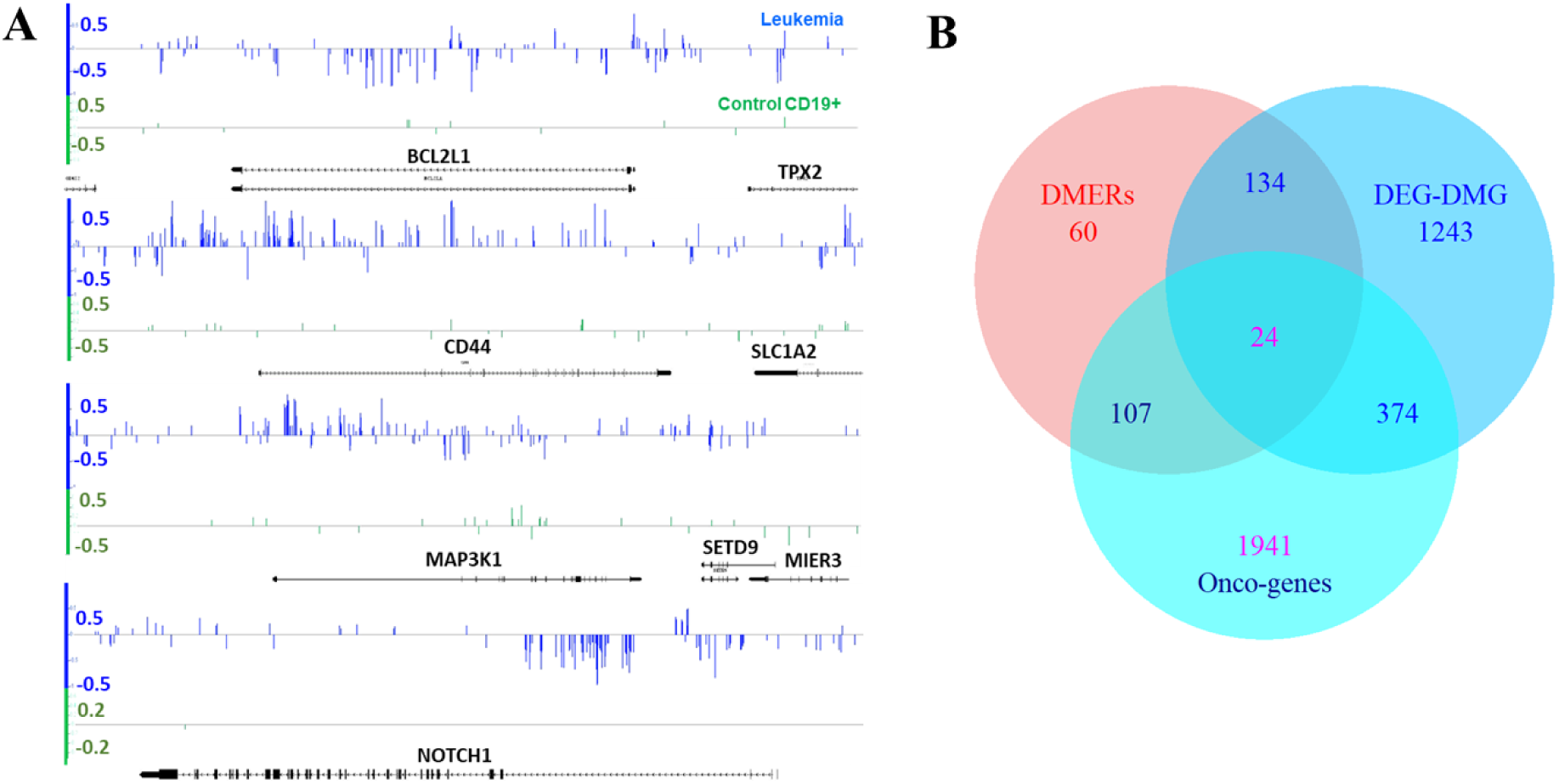
DEG-DMGs reported as cancer related gene (oncogene) lists. **A**, Group mean of methylation level differences at each cytosine identified differentially methylated genes (DMGs). BCL2L1, CD44, MAP3K1 and NOTCH1 are linked to leukemia and other types of cancers. These genes were identified as “hubs” of PPI networks (Fig. 5B and D). Irregular distribution of methylation signal, hyper- and hypo-methylated, can be viewed. Traditional DMR-based approaches fail to detect these types of variation. Methylation level differences were computed for control and treatment individuals with respect to normal CD19+ methylome from four independent blood donors used as reference. This approach provides an estimation of the natural variability of methylation changes existing in the control population. **B**, Overlapping (≥500bp) between the differentially methylated enhancer regions (DMERs) and DEGs-DMGs. Although only 51 enhancers (DMERs) are activators of reported DEGs, the DMERs overlap with 159 DEGs-DMGs regions, from which 23 are reported oncogenes (see Methods). A total of 379 DEGs-DMGs are reported oncogenes.

Three members of this pathway are found in the sub-network DMG-DEGs shown in Fig. 5D and in the DMP distribution on MAP3K1 gene-body shown in Fig. 6A. In whole, 379 identified DEG-DMGs have been reported as cancer-related genes (Fig. 6B).

### Differentially methylated enhancer regions (DMERs)

Our initial analysis was limited to the methylation signal carried on gene-body regions. As suggested in Fig. 6, gene-associated methylation signal can also be present on genomic regions upstream and downstream to genes, including transcription enhancer regions ^22^. Analysis of the methylation datasets identified 325 differentially methylated enhancer regions (DMERs). Although only 51 enhancers from the 325 identified DMERs are activators of reported DEGs (Supplementary Table S2), the list of DEG-DMG regions covered by DMERs (in at least 500bp) reach a total of 159 (Fig. 6B), from which 23 were identified oncogenes.

The top 29 genes with highest density variation of DMP number within enhancer regions are shown in Figure 7. Many of these genes have been reported to be associated with cancer development and were found in the sets of DMGs or DEGs. One example is the enhancer region influencing gene *EPIDERMAL GROWTH FACTOR-LIKE DOMAIN 7 (EGFL7*) and the micro-RNA *MIR-126*, both associated with cancer ^23,24^. As shown in Figure 7B, *MIR-126* resides within an intron of *EGFL7* and their enhancer region overlaps.

**Figure 7.**
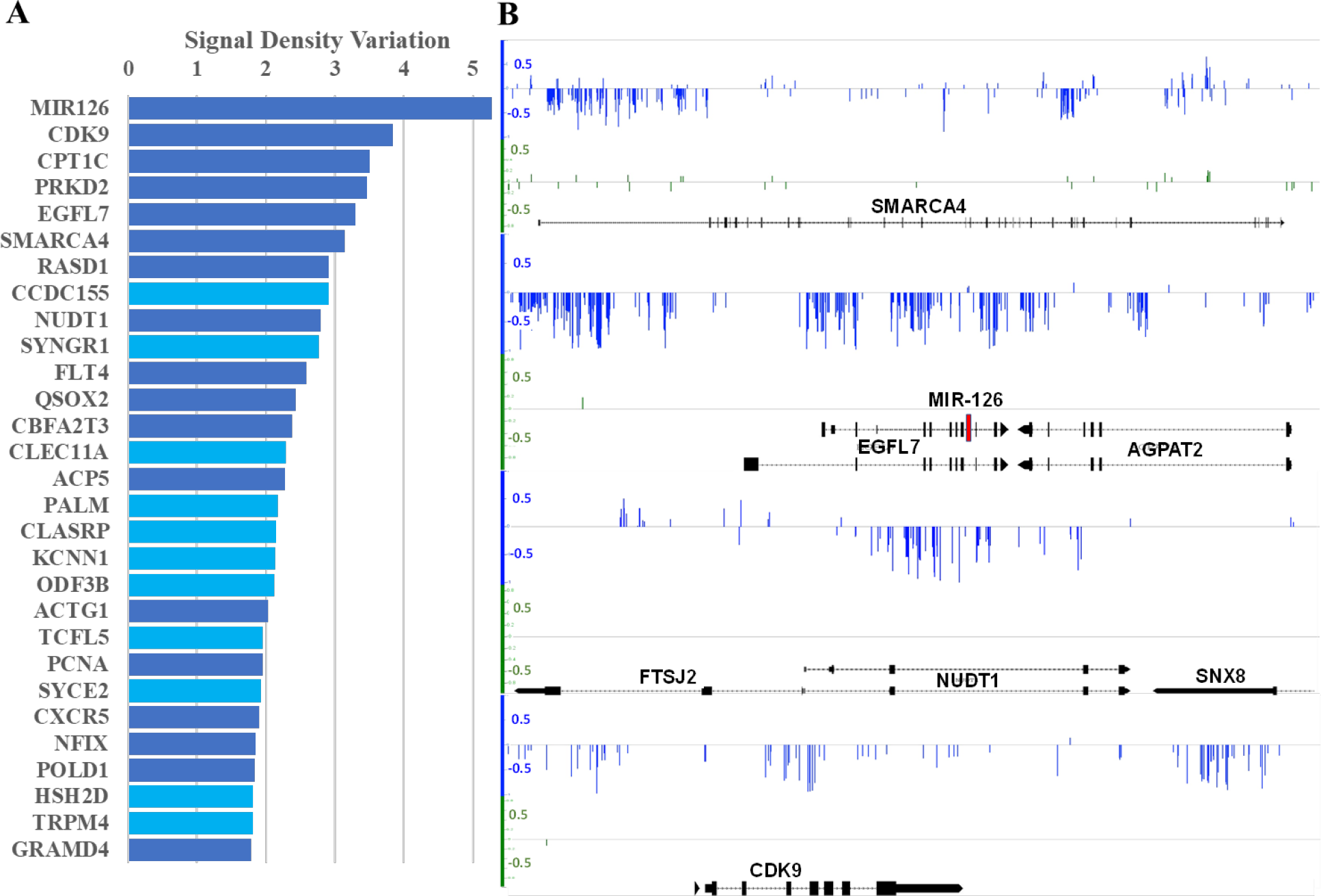
DEGs with differentially methylated enhancer region. A, Top 29 genes with the highest density variation of DMP number (> 1.7 DMPs/kb) in the enhancer region. Bars in dark blue denote genes that have been reportedly associated with cancer development. B, Group mean of methylation level differences at each cytosine identified differentially methylated enhancer regions corresponding to the genes: SMARCA4, EGRL7, MIR126, NUDT1, and CDK9.

MIR-126 modulates vascular integrity and angiogenesis, and it has been reported that MIR-126 delivered via exosomes from endothelial cells promotes anti-tumor responses ^25^. The hypomethylation pattern observed in the region spans a substantial part of gene *AGPAT2*, which was identified as a DMG and, although over-expressed in different types of cancer, was not reported as a DEG in the earlier PALL study ^26^. *AGPAT2* promotes survival and etoposide resistance of cancer cells under hypoxia ^27^.

### Association between methylation and gene expression

Results to date suggest the existence of an association, or at least statistical inter-dependence, between methylation and gene expression. To investigate this association, density variations of the methylation signal were quantitatively expressed by different measurements: density of methylation level difference |Δ*p*_*density*_|, density of total variation difference |Δ*TV*_*density*_|, and |Δ*HD*_*density*_| (see method section for variable descriptions). Gene expression was shown as absolute value of the logarithm base 2 of fold change, |*log*_2_*FC*|.

The association between methylation and gene expression for the current study of patients with PALL is shown in Supplementary Fig. S7. This association is not only corroborated by a highly significant Spearman’s rank correlation rho (*p-value* lesser than 0.001, Supplementary Fig. S7), but also by two-dimensional kernel estimation (2D-KDE) and Farlie-Gumbel-Morgenstern (FGM) copula of joint probability distributions for each annotated pair of variables in the coordinate axes from the contour-plot plane (Supplementary Fig. S7).

Results indicate that methylation and gene expression show positive dependence. Roughly speaking, a bivariate distribution is considered to have a specific positive dependence property if larger values of either random variable are probabilistically associated with larger values of the other random variable ^28^. According to Lai^29^, the FGM copulas shown in Supplementary Fig. S7 indicate CDM and gene expression to be *positively quadrant dependent* and *positively regression dependent*. In other words, if *X* is the density of methylation level difference, the regression *E*(*Y*|*X* = *x*) is linear in *x* ^29^. Thus, the regression of the conditional expected value of gene expression with respect to density variations of methylation signal *X* is linear in *x* (possible values of *X*). This linear trend is noticed with high joint probability in the outlined contour-plot red regions (Supplementary Fig. S7).

### PC-score of DEG-DMGs

The identification of genes playing fundamental roles in cancer progression is limited by the availability of protein-protein interaction information in a database (STRING, in the current case). Consequently, results could be mostly populated with genes from network-associated diseases. To circumvent these possible limitations, principal component analysis (PCA) was applied to score genes according to their discriminatory power to discern the disease state from healthy. PCA was performed on the set of individuals, representing each in the 1775-dimensional space DEGs-DMGs, where each gene was represented by the density of an information divergence on gene-body, which provides a normalized measurement of the intensity of the methylation signal. Two PC-scores were derived from two information divergences: 1) absolute difference of methylation levels and 2) Hellinger divergence.

The first principal component (PC1) was used to build the PC-scores for DMGs, since it carried 85% of the whole sample variance with eigenvalues greater than 1 (Guttman-Kaiser criterion ^30^, see Methods). A list of the first 12 genes with top PPI-network PC-scores is presented in Table 1, indicating genes associated with cancer development and further confirming that regardless of approach followed, genes involved with cancer origin and progression are DEG-DMGs.

**Table 1.**
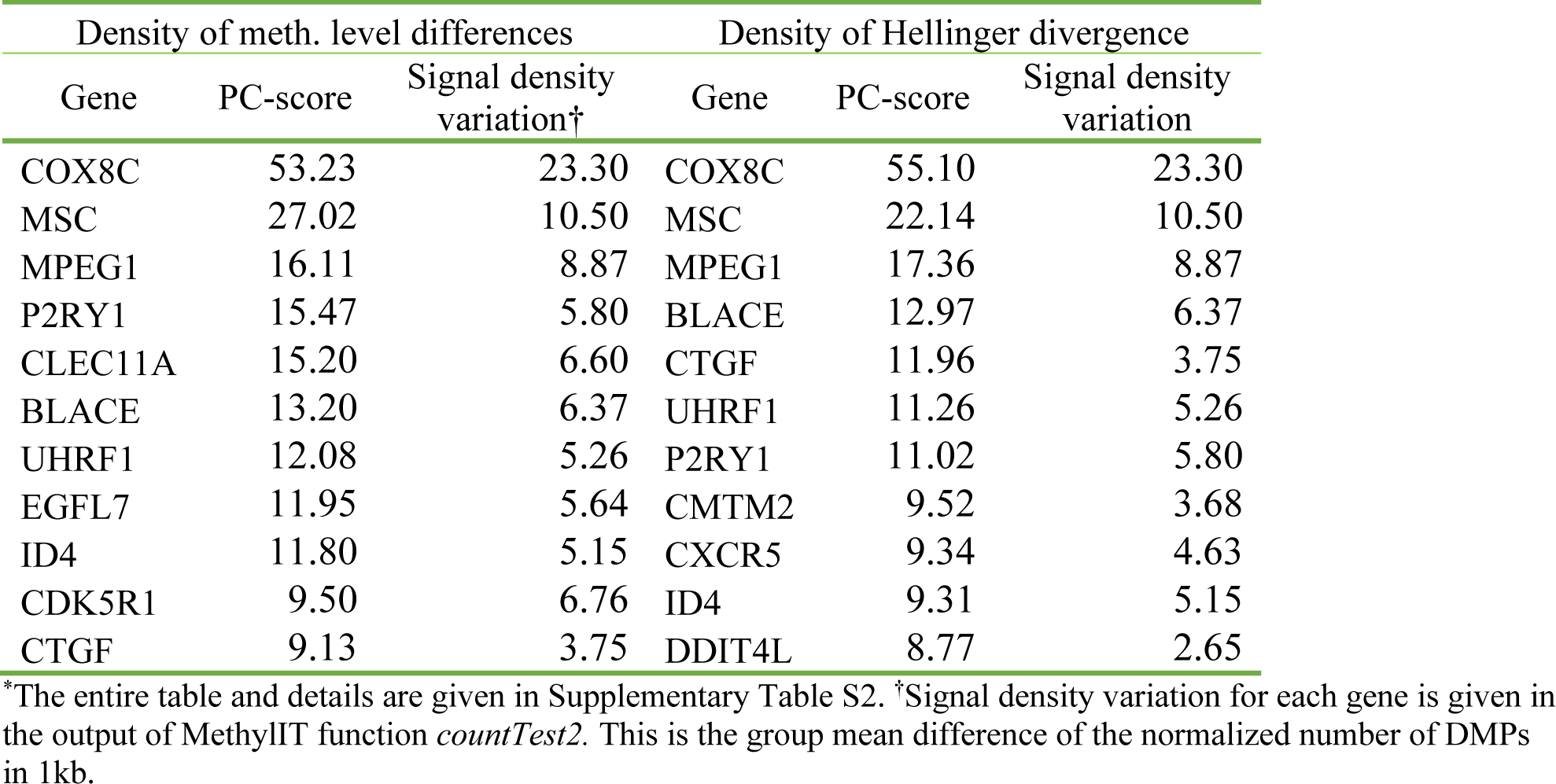
First 12 genes with the top PC-score based on density of methylation level differences and density of Hellinger divergences*.

| Density of meth. level differences |  |  | Density of Hellinger divergence |  |  |
| --- | --- | --- | --- | --- | --- |
| Gene | PC-score | Signal density variation<sup>†</sup> | Gene | PC-score | Signal density variation |
| COX8C | 53.23 | 23.30 | COX8C | 55.10 | 23.30 |
| MSC | 27.02 | 10.50 | MSC | 22.14 | 10.50 |
| MPEG1 | 16.11 | 8.87 | MPEG1 | 17.36 | 8.87 |
| P2RY1 | 15.47 | 5.80 | BLACE | 12.97 | 6.37 |
| CLEC11A | 15.20 | 6.60 | CTGF | 11.96 | 3.75 |
| BLACE | 13.20 | 6.37 | UHRF1 | 11.26 | 5.26 |
| UHRF1 | 12.08 | 5.26 | P2RY1 | 11.02 | 5.80 |
| EGFL7 | 11.95 | 5.64 | CMTM2 | 9.52 | 3.68 |
| ID4 | 11.80 | 5.15 | CXCR5 | 9.34 | 4.63 |
| CDK5R1 | 9.50 | 6.76 | ID4 | 9.31 | 5.15 |
| CTGF | 9.13 | 3.75 | DDIT4L | 8.77 | 2.65 |

\*The entire table and details are given in Supplementary Table S2. <sup>†</sup>Signal density variation for each gene is given in the output of MethyLIT function *countTest2*. This is the group mean difference of the normalized number of DMPs in 1kb.

## Discussion

Data from this study reflect non-random methylation repatterning that targets gene networks reportedly associated with cancer development and risk. The majority of DNA methylation changes fall within intergenic regions of the genome, and only 4795 (including non-coding) of the 57241 annotated human genes were identified as DMGs. This result suggests that in patients with PALL, the methylation machinery may selectively target specific genes. The methylation signal is observed not only within gene-body regions of DMGs, but also (and frequently with high intensity) in upstream and downstream domains.

Network analysis of DMGs identified several KEGG pathways and genes associated with cancer. Relevant genes were identified as network hubs and grouped into clusters of network hubs carrying the highest network centrality measurements (Fig. 1 and 5). Presumably, disruption of a network hub by altering the gene, or others that regulate the hub, could alter the entire gene network ^14,31^. Thus, identification of hubs offers candidate targets in the search for potential biomarkers. The strong linearity trends observed in pairwise regression between the centrality measurements (Fig. 2) in the main hub cluster (Fig. 1A) suggests that genes from the cluster are non-randomly targeted by the action of methylation regulatory machinery during PALL development ^14^.

Clusters of hubs integrating PPI subnetworks comprise the backbone of a network. The essentiality of gene hubs in preserving the integrity of the interacting network is quantitatively expressed in network centrality statistics. For sub-networks of hubs (Fig. 1 and 5), higher centrality values and linear relationships between the centrality statistics of the network hubs reflects a higher number of reported biologically meaningful associations between the hubs and the other genes on the sub-network and the main network (Fig. 2).

Strong correspondence was seen in the network enrichment analyses derived from PPI networks in DMGs and DEGs (Fig. 3), supporting the non-random nature of methylation signals within protein-coding regions in signaling pathways linked to cancer development. Although not all DEGs are detected as DMGs and vice versa, massive overlap of enriched KEGG pathways (Fig. 3) suggest a coordinated response of methylation and gene-expression machineries. This *in concert* regulatory response was statistically supported by Lin’s concordance correlation coefficient and Kendall coefficient of concordance.

An example of coordinated regulatory response of methylation and gene expression is seen in the case of the *EGFR* gene, identified as a hub in the DMG network (Fig. 1). *EFGR* is a tyrosine kinase that regulates autophagy via the PI3K/AKT1/mTOR, RAS/MAPK1/3 (enriched pathways shown in Fig. 3A and B, and in Fig. 5A and E), and STAT3 signaling pathways ^32,33^. Although EGFR was not a reported DEG, its activators, *EPIDERMAL GROWTH FACTOR* (*EGF*, Fig. 5B) and *EGFL7* were identified as both DMGs and DEGs. *EGFL7* is reported to be a key factor for the regulation of the *EGFR* signaling pathway ^34^. Additionally, *EGFL7* is a secreted angiogenic factor that can result in pathologic angiogenesis and enhance tumor migration and invasion via the *NOTCH* signaling pathway ^23^ (a pathway enriched in the PPI-DMG network). The *NOTCH* pathway is a conserved intercellular signaling pathway that regulates interactions between physically adjacent cells. In the set of patients with PALL, *NOTCH1* is reported as a DEG and DMG (Fig. 1A and 5B).

Another example of the gene network architecture of leukemia emerges by tracking up-and downstream interconnections of genes *PIK3CG* (DEG-DMG) and *PIK3CD* (a DMG network hub, Fig. 1) from the PI3K/AKT signaling pathway (enriched in the set of DEG-DMGs, Fig. 5). Phosphatidylinositol-4,5-bisphosphate 3-kinase (*PI3K*) is a critical node in the B-cell receptor (BCR, a DEG-DMG) signaling pathway and its isoforms, *PIK3CD* and *PIK3CG* are involved in B-cell malignancy ^35^. Crosslinking CD19 with the BCR augments PI3K activation, and VAV proteins, VAV1 (DMG), VAV2 (DEG-DMG), and VAV (DEG-DMG) also contributes to PI3K activation downstream of BCR and related receptors ^36^. BCR and its downstream signaling pathways, including Ras/Raf/MAPK, JAK/STAT3, and PI3K/AKT (all enriched in PALL patients, Fig. 3 and 5), play important roles in malignant transformation of leukemia ^37^.

Our analysis also considered gene regulatory domains upstream and downstream to gene-body regions and, in particular, enhancer regions. The set of genes targeted by DMERs does not integrate to a PPI network, but is found in signaling pathways or regulators from them. As in the previous analyses, enhancer methylation repatterning identifies genes known to be involved in cancer development (Fig. 6B). For example, *SMARCA4* (Fig. 7) encodes an ATPase of the chromatin remodeling SWI/SNF complexes frequently found upregulated in tumors ^38^ and represents a DEG-DMG in patients with PALL. The product of this gene can bind BRCA1 (DEG-DMG) ^39^ and also regulates the expression of the tumorigenic protein CD44 (DEG-DMG) ^40^.

PPI networks are only models to identify highly interconnected players from the subjacent web architecture of genes involved in a specific biological process. Thus, results from the application of more than one network model can complement, and different network models do not necessarily overlap 100% with the set of enriched pathways. Deriving subsets of the DEG-DMG dataset by applying MCODE clustering increased confidence over previous results.

The integrative analyses of DMGs, DEGs and the networks derived from them, as well as DMERs (graphically summarized in Fig. 1 to 7), provided consistent indication of a web of interacting genes in cancer development and an association between gene methylation repatterning and gene expression changes. This association was supported by Spearman’s rank correlation rho and the bivariate FGM copula (Supplementary Fig. S7), which implies a linear dependence for expected values of gene expression changes in methylation changes for the set of DEG-DMGs.

Our analysis uncovered a *stochastic deterministic dependence* relationship, where larger values of gene expression changes are probabilistically associated with larger values of methylation changes (in the whole set of 1772 DEG-DMGs). Within the set of DEG-DMGs, observed changes in gene expression were not statistically independent of the methylation changes, showing association with a significant linear trend (Supplementary Fig. S7). This result may be indication that the relationship between gene methylation repatterning and altered gene expression would be present at lower density methylation levels. Such a relationship can be overlooked with over-stringent filtering of methylome data. Three analytical approaches assist in discovering this association: *i*) signal detection for DMP identification, *ii*) GLM-based group comparison for DMG identification, and *iii*) copula modeling of stochastic dependence.

Our results demonstrate the potential of integrative network analysis of DMGs and DEGs for the identification of biologically relevant methylation biomarkers. Numerous clusters of interacting genes are detected in the sub-networks of hubs from PPI networks of DMGs and DEGs, a few of which are described here. More detailed analysis of these data has allowed us to propose three factors likely to be important to biomarker identification. A potential biomarker must 1) be a DMG or a DEG-DMG with one or more well defined differential methylation pattern(s) on gene-body, upstream or downstream of the gene-body; 2) integrate one or more gene pathways that are biologically relevant for leukemia and, simultaneously, show enrichment in the PPI networks of DMGs and DEGs, and 3) represent a hub or be biologically connected to a relevant hub. Genes holding to these guidelines integrate the subnetworks of hubs shown in Figs. 1B and 4C-D, and the list of potential biomarkers can be extended using the information provided in the Supplementary Tables S1 and S2.

Intersection of the identified networks with available data from independent studies of cancer further supports the potential of our approach for identifying robust disease biomarkers. However, while intersection of methylome and gene expression data with cancer-relevant gene networks is compelling, we cannot eliminate the possibility that these outcomes may be influenced by the relative abundance of cancer-related networks within the various databases currently available. To help circumvent this limitation, we proposed ranking the DEG-DMGs based on their discriminatory power to discern disease state from healthy.

Potential biomarkers can be scored with the application of PCA (Table 1 and Supplementary Table S2). In this study, the first PC was sufficient to build a PC-score of DEG-DMGs based on gene-body methylation signal intensity. PC-scores identify cancer-related genes not identified by the PPI network approach, although not all relevant genes were identifiable, e.g., *NOTCH1*. Within a long gene like *NOTCH1*, the non-homogenous distribution of gene body methylation signal (Fig. 6A) can result in apparently low density methylation signal globally, even when signal is high locally. Nevertheless, PC-score provides an acceptable complement to the PPI network approach. Results obtained with the approach proposed here support its application to the identification of reliable and stable biomarkers for cancer diagnosis and prognosis. Lists of genes relevant as biomarker candidates for leukemia (several of which have already been proposed as biomarkers by others) are provided in the Supplementary Tables online.

## Materials and Methods

### Methylation and gene expression datasets

The datasets of genome-wide methylated and unmethylated read counts (for each cytosine site) from normal CD19+ blood cell donor (NB) and from patients with pediatric acute lymphoblastic leukemia (PALL) where downloaded from the Gene Expression Omnibus (GEO) database. DMPs were estimated for control (NB, GEO accession: GSM1978783 to GSM1978786) and for patients (ALL cells, GEO accession number GSM1978759 to GSM1978761) relative to a reference group of four independent normal CD19+ blood cell donor (GEO accession: GSM1978787 to GSM1978790). The datasets of DEGs from the group of patients with PALL were taken from the Supplementary information provided in the previous study ^7^.A list of 2,579 cancer-related genes compiled by Bushman Lab (http://www.bushmanlab.org/links/genelists) was used to identify DEG-DMGs oncogenes.

### Methylation analysis

Methylation analysis was performed by using our home pipeline Methyl-IT version 0.3.1 (a R package available at https://git.psu.edu/genomath/MethylIT). Estimation of differentiallymethylated positions (DMPs) is consistent with the classical approach using Fisher’s exact test except for a further application of signal detection (see examples of methylation analysis with MethylIT at https://github.com/genomaths/MethylIT). Need for the application of signal detection in cancer research was pointed out decades ago ^41^. Here, application of signal detection was performed according to standard practice in current implementations of clinical diagnostic tests ^42–44^. That is, optimal cutoff values of the methylation signal were estimated on the receiver operating characteristic curves (ROCs) based on ‘Youden Index’^42^ and applied to identify DMPs. The decision of whether a DMP is detected by Fisher’s exact test (or any other statistical test implemented in Methyl-IT) is based on optimal cutoff value ^43^.

#### Estimation of differentially methylated regions (DMRs)

The regression analysis of the generalized linear model (GLMs) with logarithmic link, implemented in MethylIT function *countTest*, was applied to test the difference between groups of DMP numbers/counts at specified genomic regions, regardless of direction of methylation change. Here, the concept of DMR is generalized and it is not limited to any specific genomic region found with specific clustering algorithm. It can be applied to any naturally or algorithmically defined genomic region. For example, an exon region identified statistically to be differentially methylated by using GML is a DMR. In particular, a DMR spanning a whole gene-body region shall be called a DMG. DMGs were estimated from group comparisons for the number of DMPs on gene-body regions between control (CD19+ blood cell donor) and ALL cells based on generalized linear regression.

The fitting algorithmic approaches provided by *glm* and *glm*.*nb* functions from the R packages *stat* and MASS were used for Poisson (PR), Quasi-Poisson (QPR) and Negative Binomial (NBR) linear regression analyses, respectively. These algorithms are implemented in the Methyl-IT function *countTest* and *countTest2*, which only differ in the way to estimate the weights used in the GLM with NBR. The following *countTest* parameters were used: minimum DMP count per individual (8 DMPs), test *P-value* from a likelihood ratio test (test = “LRT”) and P-value adjustment method (Benjamini & Hochberg^45^), cut off for *P-value* (α = 0.05), and *Log2Fold* Change for group DMP number mean difference >1.

The methylation analysis of genomic regions to identify differentially methylated enhancer regions (DMERs) was performed on a set of enhancers reported by Hnisz et al. ^46^. Usually, the size of the genomic region covered by an enhancer varies depending on the tissue type. In our current case, for each enhancer we analyzed the maximum region spanning all reported sizes for different tissues.

### Network analysis

Protein-protein interaction (PPI) networks were built with STRING app of Cytoscape ^11,12^. Network analysis were conducted in Cytoscape. When the number of genes exceeded l00 for network analysis, biologically meaningful web connections were difficult to visualize. Biologically relevant subsets of genes were obtained from the whole set of genes (DMGs, DEG, or DEGs-DMGs) by using the R packages NBEA and NEAT ^9,10^. Alternatively, Cytoscape app MCODE was then used for subsetting an entire network ^47^. PPI subnetworks from four network modules identified with MCODE are shown. MCODE parameters for degree cutoff: 10, node density cutoff: 0.01, node score cutoff: 0.2, K-score 10, and max. depth: 100. K-mean clustering algorithm was applied to each subnetwork to obtain subnetworks of hubs using the following node attributes for clustering: *betweenness-centrality, degree, closeness-centrality*, and *clustering coefficient*.

Network hubs were identified based on *betweenness-centrality* and *node degree*, where size of each node (in PPI network) is proportional to its value of *betweenness-centrality* and label font size is proportional to its *node degree*. Network enrichment analysis in KEGG pathways follows each graphic subnetwork.

### Concordance test for DEG and DMG enrichments on KEGG pathways

The concordance between DEG and DMG enrichments on KEGG pathways, derived from the PPI network via STRING app in Cytoscape, was evaluated with the application of the Lin’s concordance correlation coefficient (*ρ*_*CC*_) and Kendall coefficient of concordance (*ρ*_*KC*_). The R package *agRee* was used for the bootstrap Bayesian estimation of *ρ*_*CC*_ point value and confidence interval ^48^; while the R package *vegan* was used to compute *ρ*_*KC*_through a permutation test ^49^.

To perform the concordance test, a score was assigned to each enriched KEGG pathway from DEGs and DMGs based on the *number of genes in the pathway* and on its corresponding *statistical signification* based on its FDR *p*-value. Only pathways with FDR *p*-value lesser than 0.0004 were considered. A new variable, statistical signification (*sig*) was defined according with the scale: *sig* = 1, 2, 3, for *p*-values in the intervals (10^−5^, 10^−4^), (10^−6,^ 10^−5^), and (0, 10^−6^), respectively. The valor of *sig* = 0 was assigned to pathways not enriched in one of the group, DEGs or DMGs. For example, *Phosphatidylinositol signaling system* was not enriched in the set of PPI-DMGs and, consequently *sig*_*DMG*_ = 0, but it was enriched in the set of PPI-DEGs with *sig*_*DMG*_ = 3. Then, a new variable, named *pathway score* was defined according to the formula:

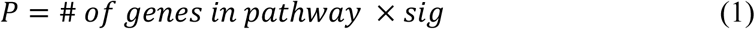

We would use the notation 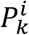 to indicate that the rating was performed for pathway *i* identified on the gene set *k* (*k =DMGs, DEGs*). That is, the pathway score *P* not only carries information on how many genes are found on each pathway but also information on the enrichment statistical signification. The estimated values of 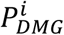 and 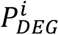 for each enriched pathway *i* (from DEGs and DMGs sets, respectively) were used in the concordance tests and in the Bland-Altman plot (Fig. 3E).

### Stochastic association between methylation and gene expression

To investigate such an association, the methylation density of gene regions simultaneously identified as DEGs and DMGs were expressed in terms of different magnitudes: 1) 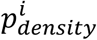, density of methylation levels (*i*: control or patients); 2) 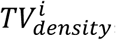, density of the difference of methylation levels between each group (control or patients) and an independent group of four healthy individuals (reference group); 3) 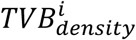, *TV* with Bayesian correction, and 4) 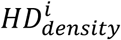, density of Hellinger divergence, where *i* denotes the group mean, control or patient. The density in 1000 bp of a variable *X* at a given gene region was defined as the sum of the magnitude *X* divided by the length of the region and multiplied by 1000. The differences of methylation densities between control and patient groups were estimated as the absolute difference of methylation levels 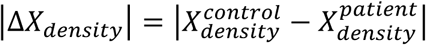, where 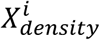 represents one of the mentioned variables. Methyl-IT R package provides all the functions to obtain all variable mentioned here (https://github.com/genomaths/MethylIT and https://github.com/genomaths/MethylIT.utils).

Spearman’s rank correlation *ρ* (rho) was estimated to evaluate the association between the pairs of variable |Δ*log*_2_*FC*| versus: |Δ*p*_*density*_|, |Δ*TV*_*density*_|, |Δ*TVD*_*density*_|, and |Δ*HD*_*density*_|. Since correlation analysis only measures the degree of dependence (mainly linear) but does not clearly discover the structure of dependence, we further investigate the structural dependence between these variables with application of Farlie-Gumbel-Morgenstern (FGM) copula. FGM copula model estimation was performed with R package copula ^50^.

### Principal component analysis (PCA)

PCA is standard statistical procedure to reduce data dimensionality, to represent the set of DMGs by new orthogonal (uncorrelated) variables, the principal components (PCs) ^51^, and to identify the variables with the main contribution to the PCs carrying most the sample variance. Herein, a PC-based score (PC-score) was built by ranking the DEG-DMGs based on its discriminatory power to discern between the disease state and healthy. Each individual was represented as vector of the 1775-dimensional space of DEG-DMGs. Two PC-scores were estimated: the first based on the density of Hellinger divergence on the gene-body and the second one based on the density of the absolute value of methylation levels difference. The density of a magnitude *x* is defined as the sum of *x* at each DMP divided by the gene width (in base-pairs). The first principal component (PC1) was used to build a PC-based score for the DEG-DMG set, since it had an eigenvalues (variance) greater than 1 and carried more than 85% of the whole sample variance (Guttman-Kaiser criterion ^30^). The PC-score was built using the absolute values of the coefficients (loadings) in PC1 for each variable (gene). Since the sum of the squared of variable loadings over a principal component is equal to 1, the squared loadings tell us the proportion of variance of one variable explained by the given principal component. Thus, the greater is the PC-score value, the greater will be the discriminatory power carried by the gene.

The density of *HD* on the gene-body was computed with MethylIT function *getGRegionsStat* and the principal component with function *pcaLDA*, which conveniently applies the PCA calling function *prcomp* from the R package ‘*stats*’.

## Acknowledgements

We wish to thank Dr. Xiaodong Yang and Thomas Maher for valuable discussions during the development of studies. This study was supported by a grant from the Bill and Melinda Gates Foundation (OPP1088661) to S.A.M.

## Author contributions

R.S. designed experiments conducted mathematical and statistics analyses. S.M. assessed experiments and edited manuscript.

## Data availability

All the methylome datasets and software used in this work are publicly available. The MethylIT R package used in the DMP and DMG estimations, as well as several examples on how to use Methyl-IT, are available at GitHub: https://github.com/genomaths/MethylIT. The datasets supporting conclusions of this report are included within Supplementary material.

## Supplementary Figures

**Supplementary Figure S1**. Distribution of methylation changes on chromosome and gene-body. **A**, Distribution of methylation changes at DMP positions on selected chromosomes as viewed within genome browser. **B**, boxplot of the means of methylation levels on chromosomes and at genes. In all cases, patient (P) data are in blue and control (C) in green.

**Supplementary Figure S2**. PPI networks built on the subset of 285 network-related DMGs. The size of each node is proportional to its value of betweeness centrality and the label font size is proportional to its node degree. Node colors from light-green to red maps the discrete scale of logarithm base 2 of fold change in DMP number for the corresponding gene: light-green: [1, 2), cyan: [2, 3), blue: [3, 4), and red: 5 or more. **B**, a subnetwork with minor hubs (101 DMGs). **C**, a cluster (139 DMGs) integrated by two subnetworks.

**Supplementary Figure S3**. PPI subnetwork module derived with Cytoscape app MCODE from the PPI network of 1775 DEG-DMGs. Node colors from yellow to red maps the discrete scale of logarithm base 2 of fold changes in gene expression for the corresponding gene: yellow: lesser or equal to −6, …, light-green: (−2, −1], …, cyan: (2, 3], … blue: (4, 5], …, red: 10 or more.

**Supplementary Figure S4**. Sub-networks derived with K-means clustering from the subset of 285 network-related DMGs.

**Supplementary Figure S5**. Network enrichment analysis on KEGG pathways for module derived with Cytoscape app MCODE from the PPI network of 1775 DEG-DMGs..

**Supplementary Figure S6**. PPI networks on the set of 191 network related DEG-DMGs. The PPI network was built with Cytoscape 11,12 from a subset of 191 DEG-DMGs previously obtained by applying network-based enrichment analysis 51. Nodes with the same color belong to the same cluster obtained by K-means clustering.

**Supplementary Figure S7**. Association between methylation and gene expression. **A**, Spearman’s rank correlation rho between variables for absolute value of logarithm base 2 of fold change (*log*_2_*FC*) in gene expression at DEG-DMGs and the differences in methylation densities (between control and patient groups). All correlations are statistically significant (*p-value* lesser than 0.001). The variables analyzed are the absolute difference (Δ) of: *pp*_*density*_: density of methylation levels, *TV*_*density*_: density of the difference of methylation levels, *TVB*_*density*_: *TV* with Bayesian correction, and *HD*_*density*_: density of Hellinger divergence of methylation levels. **B, D**, and **F** panels show two-dimensional kernel estimations (2D-KDE) of the joint probability distribution for each annotated pair of variables in the coordinate axes from the contour-plot plane (see main text for variable description). **C** and **E** panels: Farlie-Gumbel-Morgenstern (FGM) copula joint probability distribution built from the estimation of marginals distribution (*XZ* plane: Gamma probability distribution and *YZ* plane: generalized gamma distribution). Together, panels **A** to **F** indicate that, in the current study of patients with PALL, methylation and gene expression are not statistically independent, but associated with statistically highly significant linear trend, located with high joint probability in the outlined contour-plot red regions.

## Supplementary Tables

**Supplementary Table S1:** Excel files containing Tables S1-A to S1-G.

**Supplementary Table S2:** Excel files containing Tables S2-A to S1-I.

**Supplementary File S1**. zip file containing the wig files with tracks for the group means of the differences of methylation levels between each group and the reference group (four independent normal CD19+ blood cell donor): control (four normal CD19+ blood cell donor) versus reference, and patients (ALL cells from three patients) versus reference.

