## Supplementary figures and images for "Integrative Network Analysis of Differentially Methylated and Expressed Genes for Biomarker Identification in Leukemia"

### Supplementaty Fig. S1

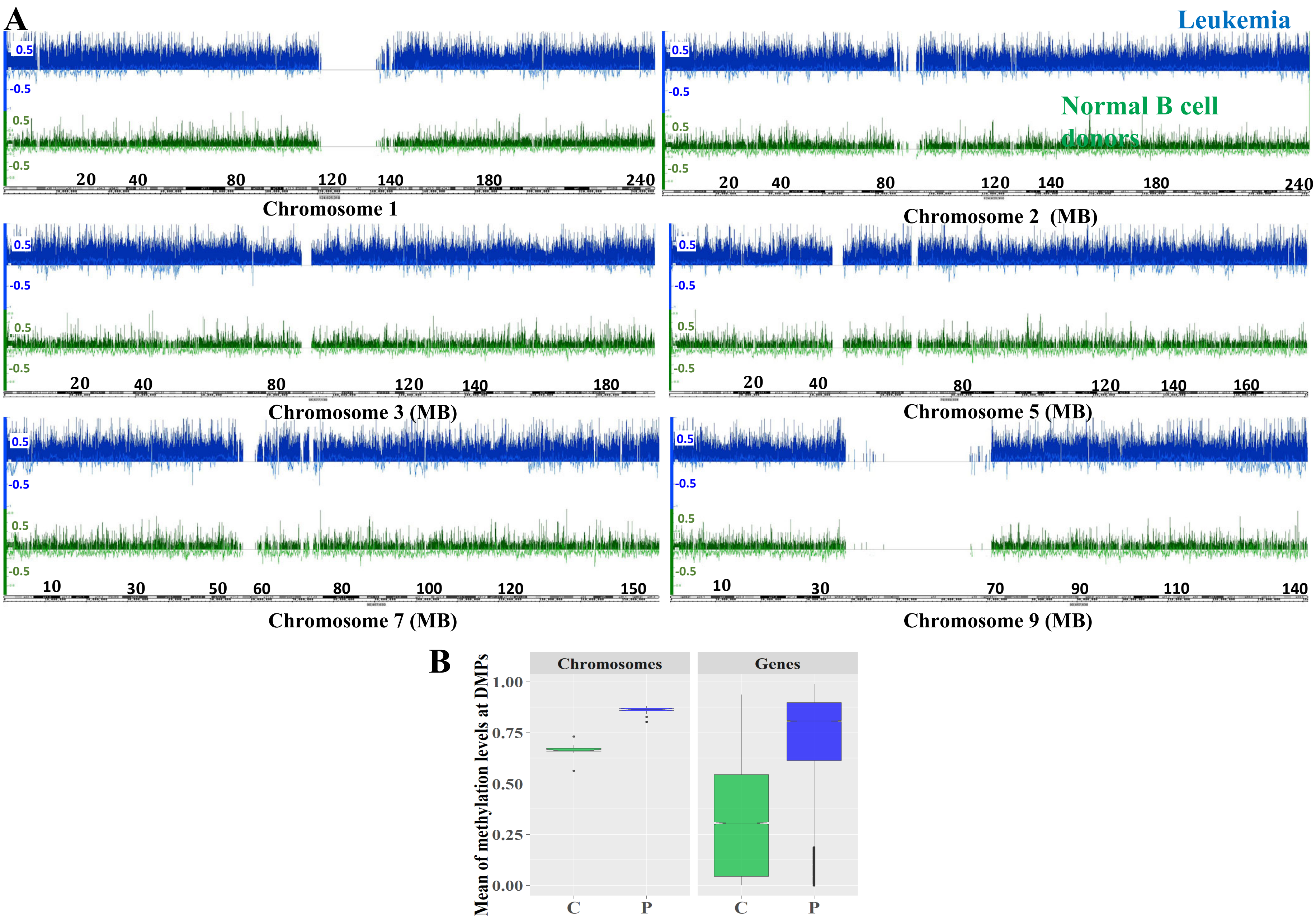

### Supplementaty Fig. S2

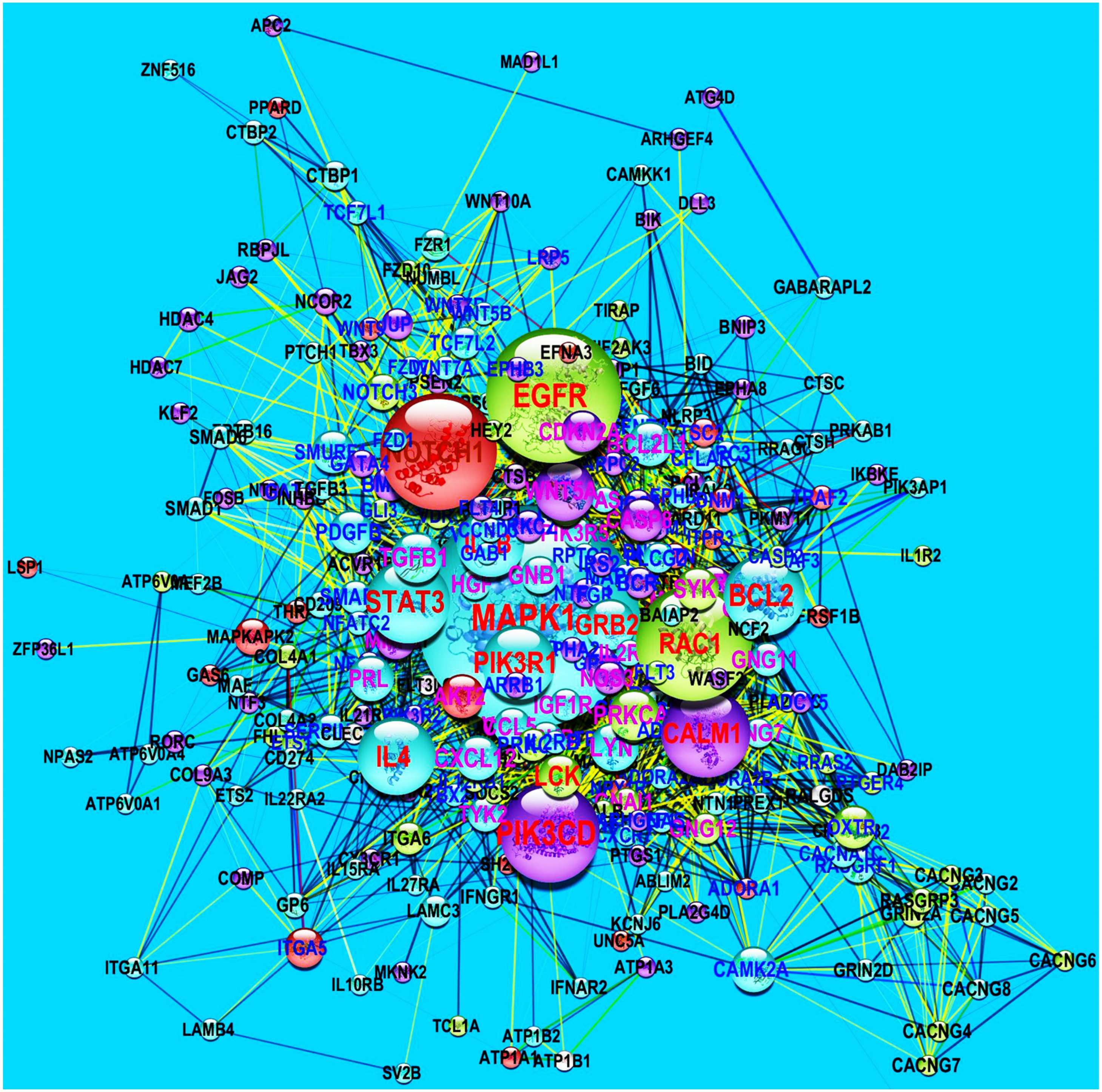

### Supplementaty Fig. S3

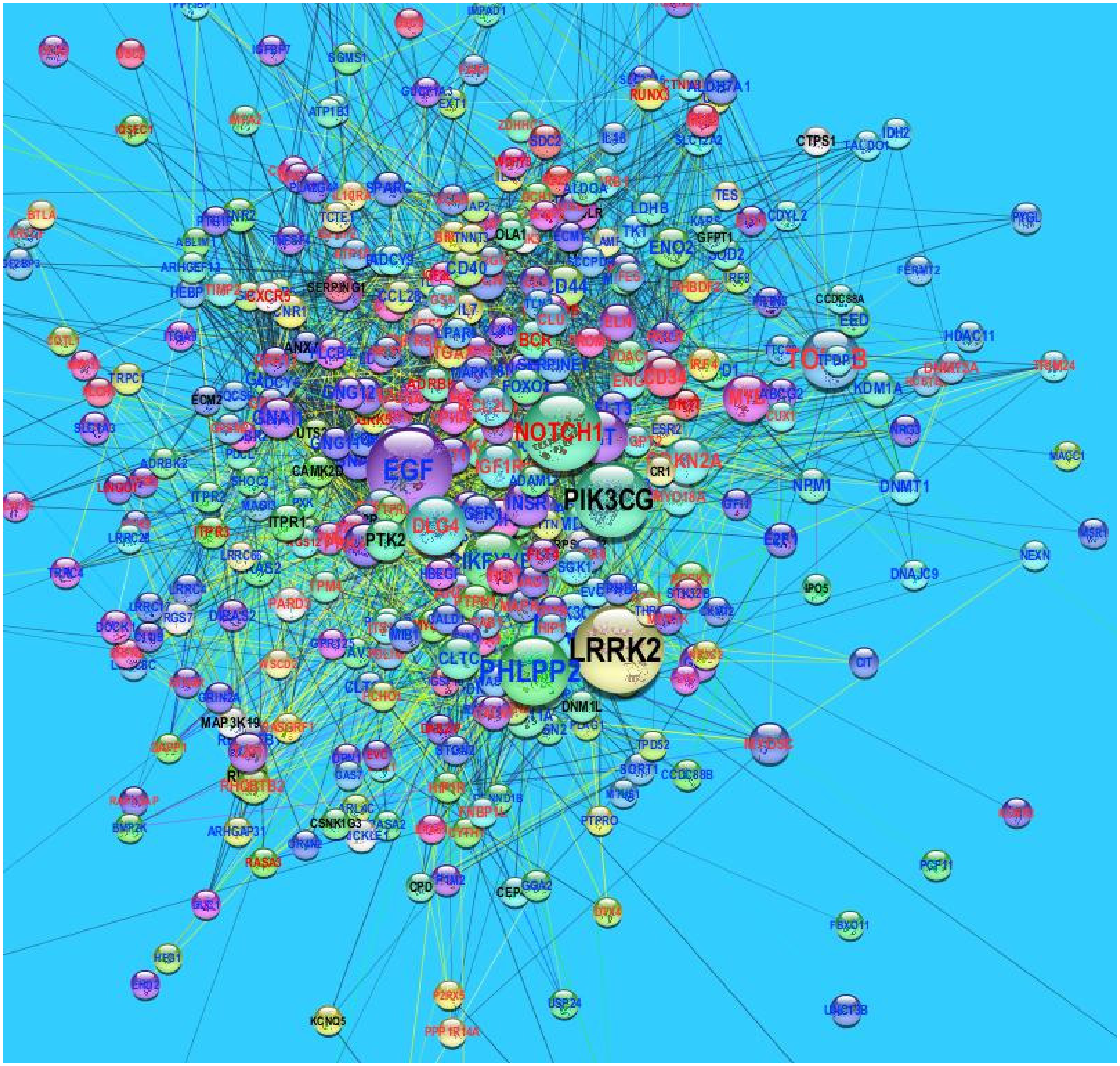

### Supplementaty Fig. S4

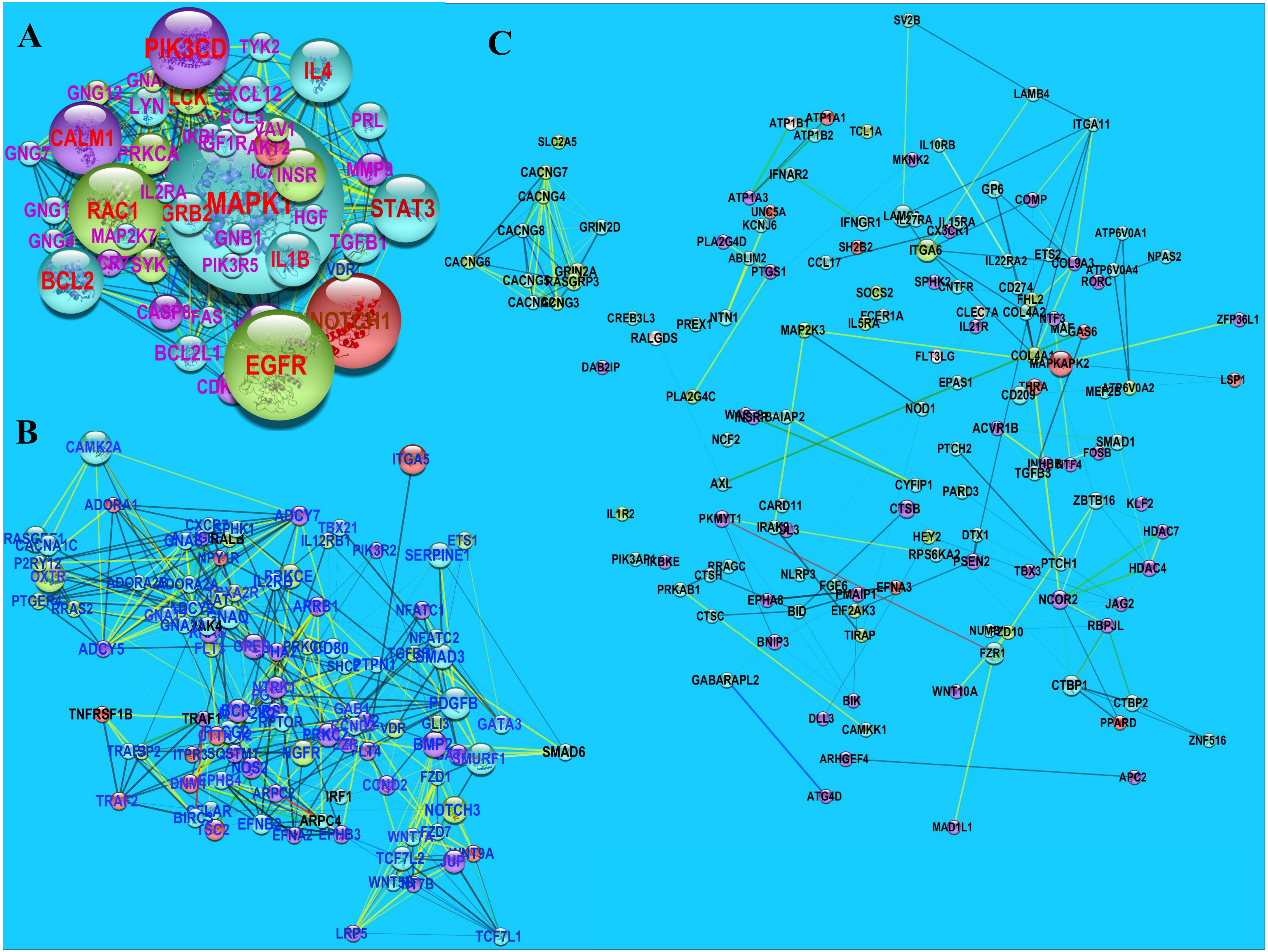

### Supplementaty Fig. S5

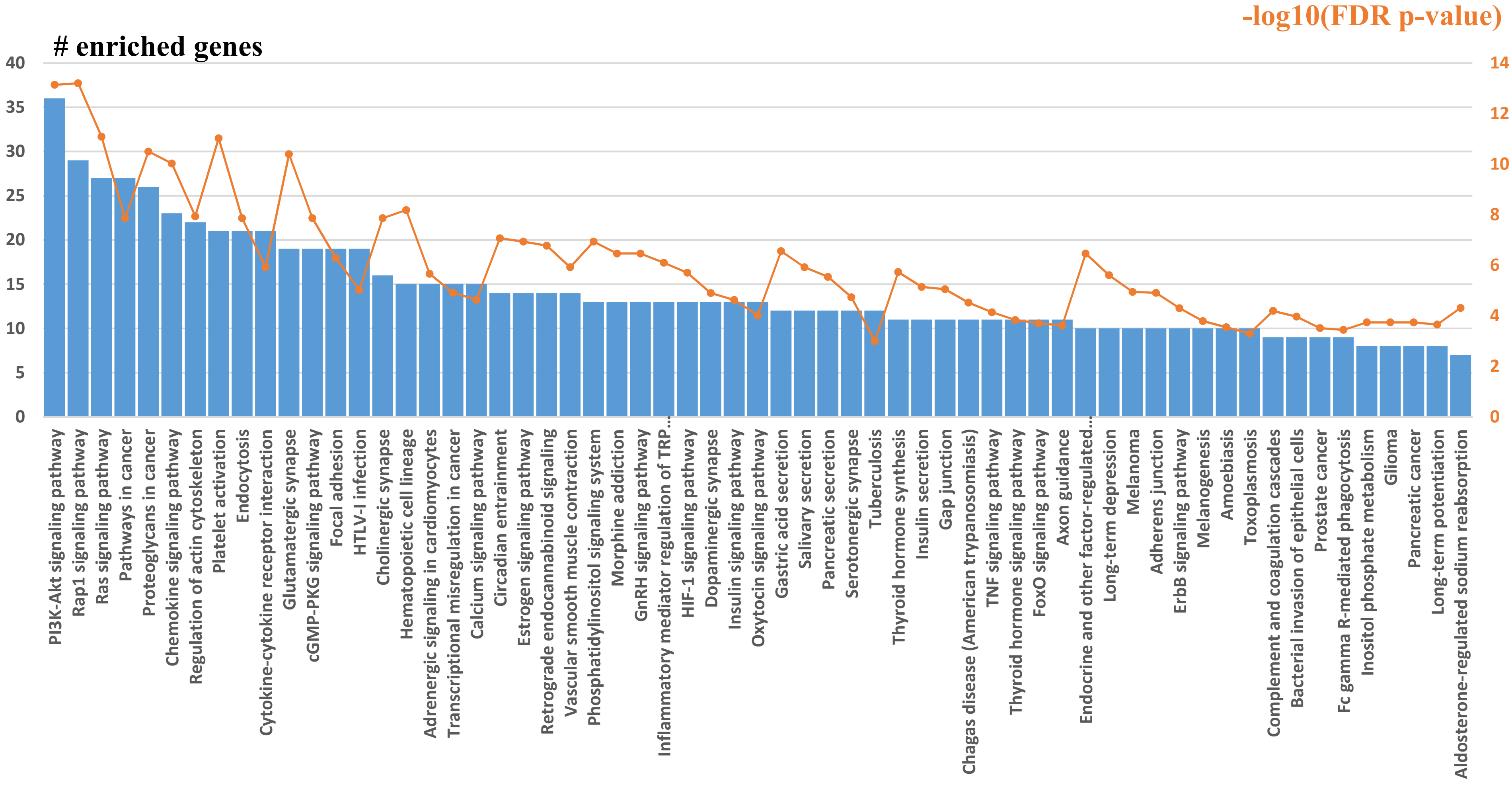

### Supplementaty Fig. S6

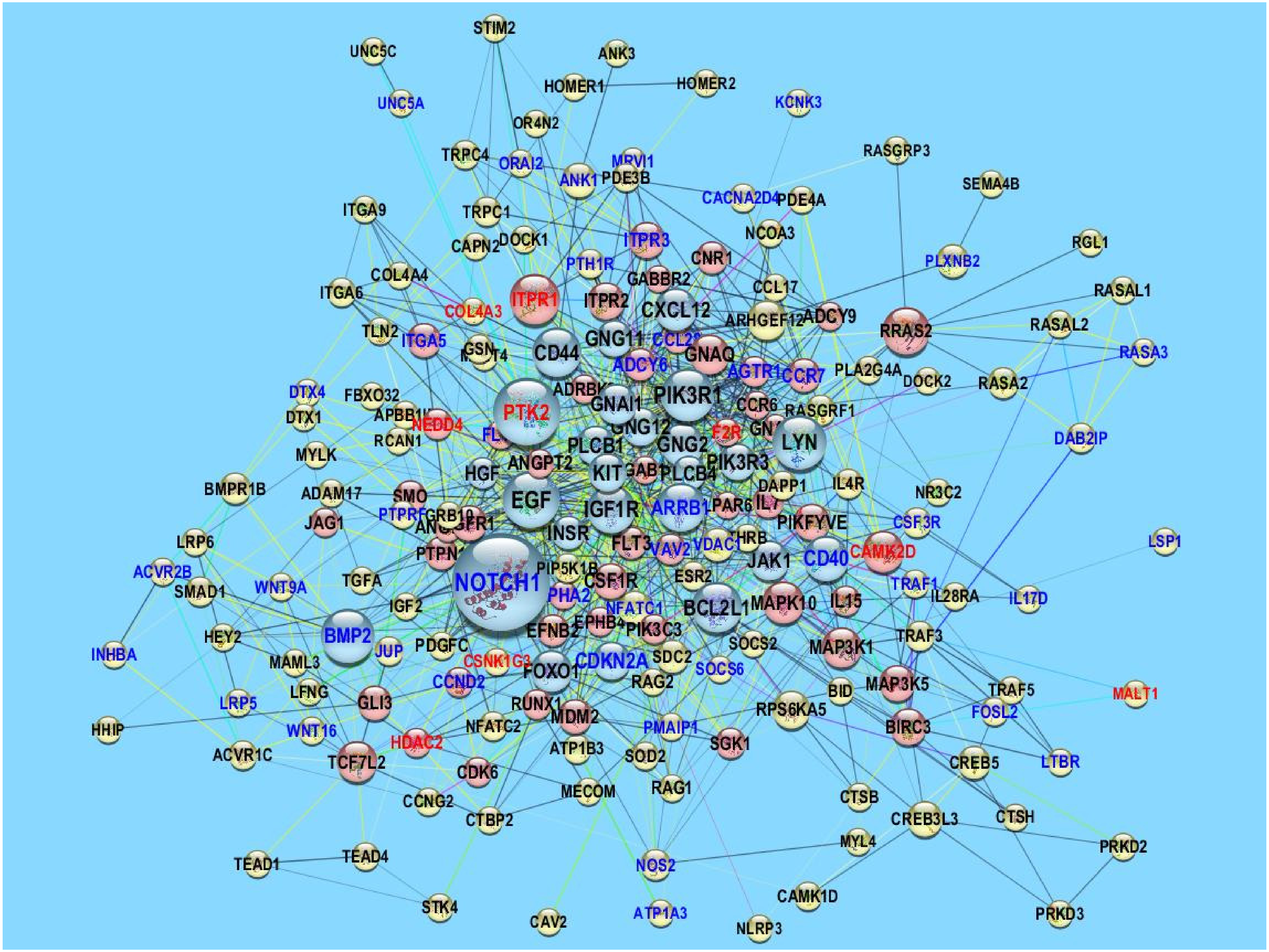

### Supplementaty Fig. S7

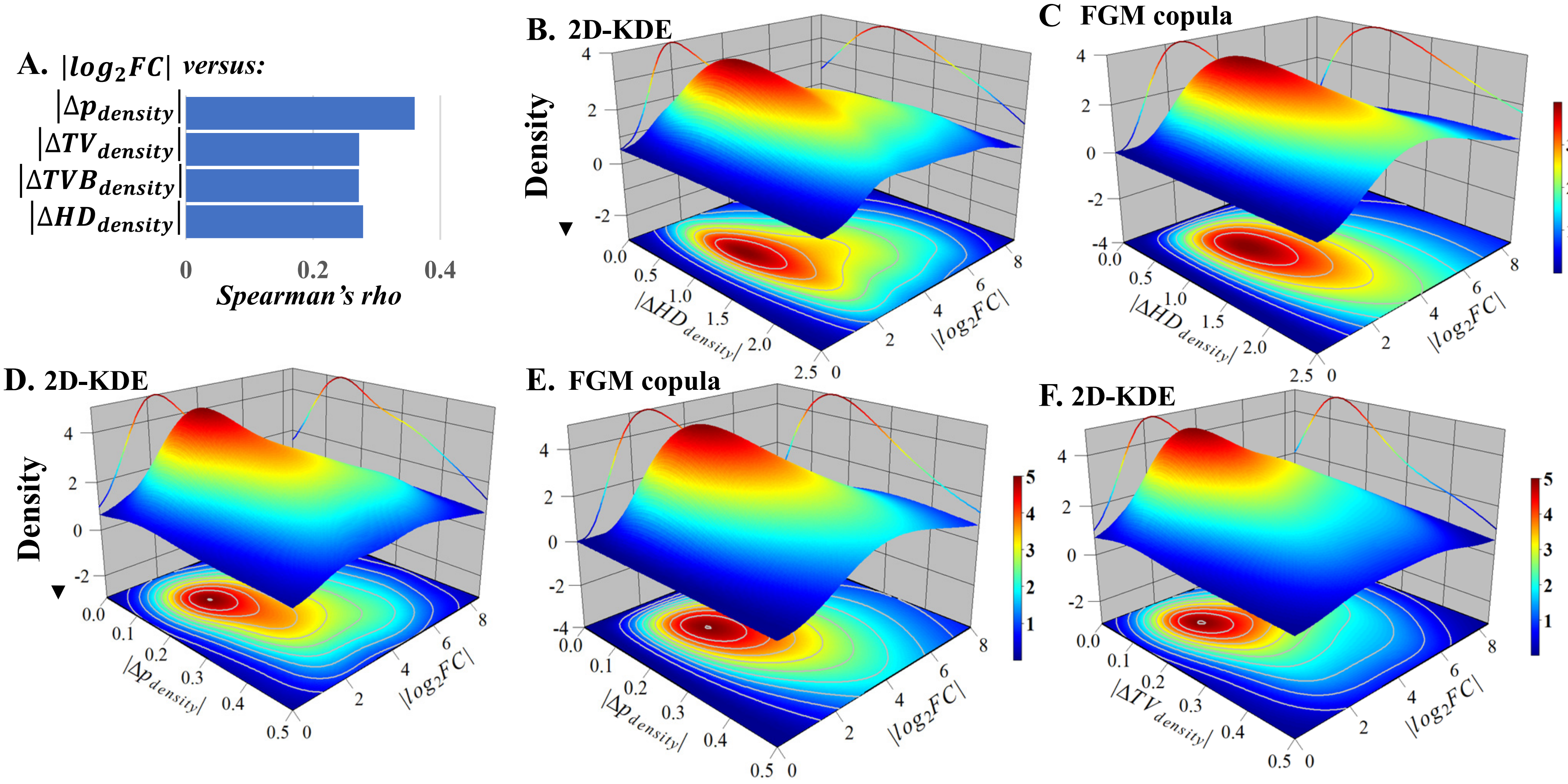
